# Pavlovian conditioned approach with different reward magnitudes reveals opposing effects of acute and chronic semaglutide treatment

**DOI:** 10.64898/2026.09.08.750149

**Authors:** Stephanie S. Desrochers, Raymond Li, Shelly B. Flagel

**Affiliations:** Michigan Neuroscience Institute, University of Michigan; College of Literature, Science, and the Arts, University of Michigan; Department of Psychiatry, University of Michigan

**Keywords:** semaglutide, motivation, incentive salience, reward-magnitude

## Abstract

**Rationale:** Glucagon-like peptide-1 receptor agonists (GLP-1RAs) have attracted interest for their effects on motivational processes beyond satiation and weight loss, including motivation for drug rewards. Reward-predictive cues can acquire motivational value and promote food- and drug-seeking, raising the possibility that GLP-1RAs affect intake by altering cue-motivated behavior. Preclinical studies report reduced food intake and responding for drug rewards following GLP-1RA treatment. However, many preclinical studies have used acute treatment; whereas, clinically, GLP1-RAs are administered as chronic treatments with dose escalation to mitigate adverse effects.

**Objectives:** Using a modified Pavlovian conditioned approach (PavCA) paradigm, we directly compared the effects of acute and chronic semaglutide treatment on conditioned approach behavior elicited by cues predicting different magnitudes of palatable food reward in male and female rats.

**Results:** Although both treatment regimens reduced body weight relative to vehicle treatment, chronic semaglutide increased, whereas acute semaglutide decreased, sign-tracking toward reward-associated cues. Rats exhibited greater sign-tracking toward cues predicting large versus small rewards, but semaglutide did not differentially alter this reward-magnitude effect. Acutely treated rats consumed fewer reward pellets and exhibited less food-cup activity during non-cue periods. These findings suggest that nonspecific behavioral suppression or malaise may have contributed to reduced responding after acute treatment.

**Conclusions:** The effects of semaglutide on cue-motivated behavior depend on the treatment regimen. Acute-treatment studies should be interpreted cautiously because adverse or nonspecific effects may contribute to apparent reductions in motivation.

## Introduction

Glucagon-like peptide-1 receptor agonists (GLP-1RAs) have become widely used for weight management and the treatment of obesity. This class of drugs was originally developed for the treatment of Type II diabetes due to their beneficial effects on blood glucose levels, but clinical studies also revealed substantial reductions in body weight (Drucker et al. 2017; Drucker 2024; Nielsen et al. 2004). These findings contributed to the development and increasing clinical use of GLP-1RAs specifically for the treatment of obesity. Peripherally, GLP-1RAs act on receptors in the gastrointestinal tract and other organs to regulate insulin release and slow gastric emptying; however, some of their weight loss effects may depend on actions in the central nervous system (Kanoski et al. 2011; Secher et al. 2014; Sisley et al. 2014; Gabery et al. 2020). Preclinical evidence suggests that, despite limited access across the blood-brain-barrier (Gabery et al. 2020; Salameh et al. 2020), GLP-1RAs may affect food intake and satiation through actions in the dorsal vagal complex, including the nucleus of the solitary tract and area postrema (Yamamoto et al. 2003; Abbott et al. 2005; Kanoski et al. 2011). Signaling from these regions may in turn influence brain areas such as the ventral tegmental area and nucleus accumbens, which contribute to dopamine-dependent reward and motivational processes (Alhadeff et al. 2012; Kooij et al. 2024; Merkel et al. 2025).

Accordingly, human studies suggest that GLP-1RAs may affect motivation beyond their impact on satiation and food intake. For example, exenatide treatment reduced the number of heavy drinking days in individuals with alcohol use disorder (Klausen et al. 2022), and semaglutide treatment reduced the number of drinks consumed on drinking days (Hendershot et al. 2025). Semaglutide treatment also reduced the number of cigarettes smoked among participants who smoke (Hendershot et al. 2025), suggesting that these drugs may alter reward-related processes underlying motivated behavior across drug classes. GLP-1RAs are also thought to influence a phenomenon known as “food noise”, in which food-associated cues can trigger intrusive thoughts about and desire for food, independent of homeostatic need (Hayashi et al. 2023). Indeed, acute exenatide administration has been shown to reduce neural responses to food images (van Bloemendaal et al. 2014) and neural activity during the anticipation of a palatable food reward (van Bloemendaal et al. 2015). These findings raise the possibility that GLP-1RAs influence motivated behavior partly by altering responses to reward-predictive cues. However, the interpretation of these effects is complicated because GLP-1RAs have clinical side effects such as nausea and vomiting (Sikirica et al. 2017), which may promote aversive learning or otherwise suppress motivated behavior. Gastrointestinal side effects are most prevalent during treatment initiation; consequently, clinical treatment typically involves gradual dose escalation to improve tolerability (Wharton et al. 2022b; Alhazmi and le Roux 2026). The effects of GLP-1RAs on motivated behavior may therefore depend on the dose, stage, and duration of treatment.

Preclinical findings support a distinction between the acute and chronic effects of GLP-1RAs on food-motivated behavior. For example, a single dose of the GLP-1RA exendin-4, administered either peripherally or directly into the NTS, decreased progressive ratio responding and conditioned place preference for sugar-pellet rewards (Dickson et al. 2012; Richard et al. 2015). In contrast, chronic semaglutide treatment induced weight loss, but also resulted in increased sucrose consumption in a 2-bottle preference test in rats (Cawthon et al. 2023), increased licking and trial initiation in a sucrose brief-access test in mice (Acosta et al. 2026), and increased progressive ratio responding for reward pellets in rats (Chang et al. 2026). Additionally, in an operant choice procedure in rats, repeated semaglutide treatment increased the preference for a palatable food reward over a cocaine infusion (Heslep et al. 2026).

Together, these findings indicate that acute and chronic GLP-1RA treatment regimens may have divergent effects on food-motivated behavior. However, these studies primarily assessed reward consumption, operant responding, or reward choice, and did not determine whether treatment regimen differentially influences the motivational value of reward-predictive cues. This distinction is important because reward-predictive cues can acquire incentive motivational value and promote reward-seeking behavior, potentially contributing to cue-triggered craving and “food noise”.

Pavlovian conditioned approach (PavCA) is a well-established paradigm for studying individual differences in the attribution of incentive motivational value to reward-predictive cues (Boakes 1977; Robinson and Flagel 2009; Meyer et al. 2012). During PavCA training, presentation of a lever-cue predicts the delivery of a reward pellet into a food-cup. Over time, subjects may develop distinct behavioral patterns reflective of different motivational processes. Goal-tracking, behavior directed toward the reward-delivery location, reflects mere learning of the cue-reward relationship, or the attribution of predictive value to the cue. Sign-tracking, behavior directed toward the reward-predictive cue, is thought to reflect the attribution of both predictive and incentive motivational value to the cue. Previous work from our laboratory showed that chronic treatment with the GLP-1RA semaglutide did not alter goal- or sign-tracking behavior during PavCA, but did increase operant responding for access to the lever-cue in a test of conditioned reinforcement (Chang et al. 2026). These findings suggest that chronic semaglutide treatment may enhance the incentive motivational value of food-reward-predictive cues, although this effect was not detectable during standard PavCA training.

In the present study, we modified the PavCA paradigm by varying the magnitude of the predicted reward (hereafter, reward-magnitude-modulated PavCA [RMM-PavCA]). Specifically, one lever-cue predicted a small reward of one pellet, whereas a second lever-cue predicted a larger reward of three pellets. This design allowed us to assess whether semaglutide altered cue-directed responding as a function of predicted reward magnitude within the same session. Additionally, we included both acute and chronic semaglutide treatment groups to allow direct comparison of treatment-regimen effects on cue-motivated behavior. We hypothesized that cues predicting different reward magnitudes would increase sensitivity to treatment-related differences by enhancing the incentive motivational value of the cue associated with the larger reward. Accordingly, based on our prior findings (Chang et al. 2026), we predicted that chronic semaglutide treatment would increase the attribution of incentive salience—indexed by sign-tracking—to cues predicting the larger reward. In contrast, we expected acute semaglutide treatment to produce a generalized reduction in motivated responding.

## Methods

All procedures were approved by the University of Michigan Institutional Animal Care and Use Committee and conformed to *The Guide for the Care and Use of Laboratory Animals*.

### Subjects

Adult male (n = 32) and female (n = 32) Sprague Dawley rats were purchased from Charles River Laboratories (Raleigh, NC and Kingston, NY). At the start of the experiment, males from Barriers R06/K98 weighed approximately 300-370 g, and females from Barriers R08/K98 weighed approximately 260-330 g. Rats were housed in same-sex pairs on ventilated racks in climate-controlled vivarium rooms and allowed to acclimate for at least 5 days prior to the start of the experiment. Throughout the experiment, rats had *ad libitum* access to standard chow (Lab Diet, Product #5LOD) and water and were maintained on a 12-hr light/dark cycle (lights on at 7:00 AM).

Rats were initially randomly assigned to either the vehicle (VEH; n = 21 males and 21 females) or chronic semaglutide (CHRONIC SEMA; n = 11 males and 11 females) treatment condition. Only one rat per cage was assigned to the CHRONIC SEMA condition, with their cage mate assigned initially to the VEH condition. Remaining cage pairs were all assigned to the VEH condition until subsequent ACUTE SEMA assignments, which were not balanced according to cage mate assignment. After PavCA session 6, the VEH group was pseudorandomly divided into VEH (n = 10 males and 10 females) and acute semaglutide (ACUTE SEMA; n = 11 males and 11 females) treatment groups, which were matched within sex for body weight and prior RMM-PavCA behavioral responding across sessions 1-6.

### Behavioral Apparatus

Experiments were conducted in 32 identical MED Associates (Fairfax, VT) behavioral testing chambers (tall rat chamber with mouse floors; ENV-007CT and ENV-005A; 29.5 x 23.6 x 27.4 cm) housed in sound attenuating cabinets equipped with ventilation fans. On one wall of each chamber, a houselight (ENV-215M) was located 1 cm below the top center of the wall. On the opposite wall, a recessed food cup equipped with an infrared head-entry detector (ENV-200R2M and ENV-254CB) was located centrally, 1.5 cm above the floor. The food cup was connected to a pellet dispenser (ENV-203/204) that delivered 45-mg banana-flavored grain pellets (Bio-Serv, Flemington, NJ; Product #F0059). Illuminable retractable levers (ENV-112CML) were located on either side of the food cup, 6 cm above the floor. The levers were calibrated to record contacts requiring approximately 15 g of force.

### Experimental Procedures

All experimental procedures occurred during the light phase between 11:00 AM and 2:00 PM. On days when behavioral testing occurred, body weight was recorded, and injections were administered after testing, approximately 24 hr before the subsequent behavioral session. One behavioral session occurred per day.

#### Semaglutide treatment

Semaglutide (AstaTech, Bristol, PA; Product AT35750) was dissolved in vehicle (44 mM sodium phosphate dibasic, 70 mM NaCl, and 0.007% Tween 20) to produce a 70 μg/mL stock solution, which was further diluted in vehicle to produce lower concentrations. Solutions of 7-70 μg/mL were administered subcutaneously (s.c.) at 1 mL/kg body weight, yielding doses of 7-70 μg/kg. Rats in the chronic semaglutide (CHRONIC SEMA) condition received once-daily escalating doses of semaglutide daily based on the protocol developed by Cawthon et al. (2023). On experimental days 1-10, rats received 7, 14, 21, 28, 35, 42, 49, 56, 63, and 70 μg/kg injections, respectively (**Fig. 1A**). They were then maintained at 70 μg/kg once daily for the remainder of the experiment (experimental days 11-20; **Fig. 1A**). Rats in the vehicle (VEH) condition received daily vehicle injections throughout the experiment (experimental days 1-20; **Fig. 1A**). Rats in the acute semaglutide (ACUTE SEMA) condition received daily vehicle injections on experimental days 1-19 and a single 70 μg/kg dose of semaglutide approximately 24 hr before the final behavioral testing session (experimental day 20; **Fig. 1A**). Thus, experimental day 20 injections determined the drug condition during PavCA session 7.

**Fig. 1.**
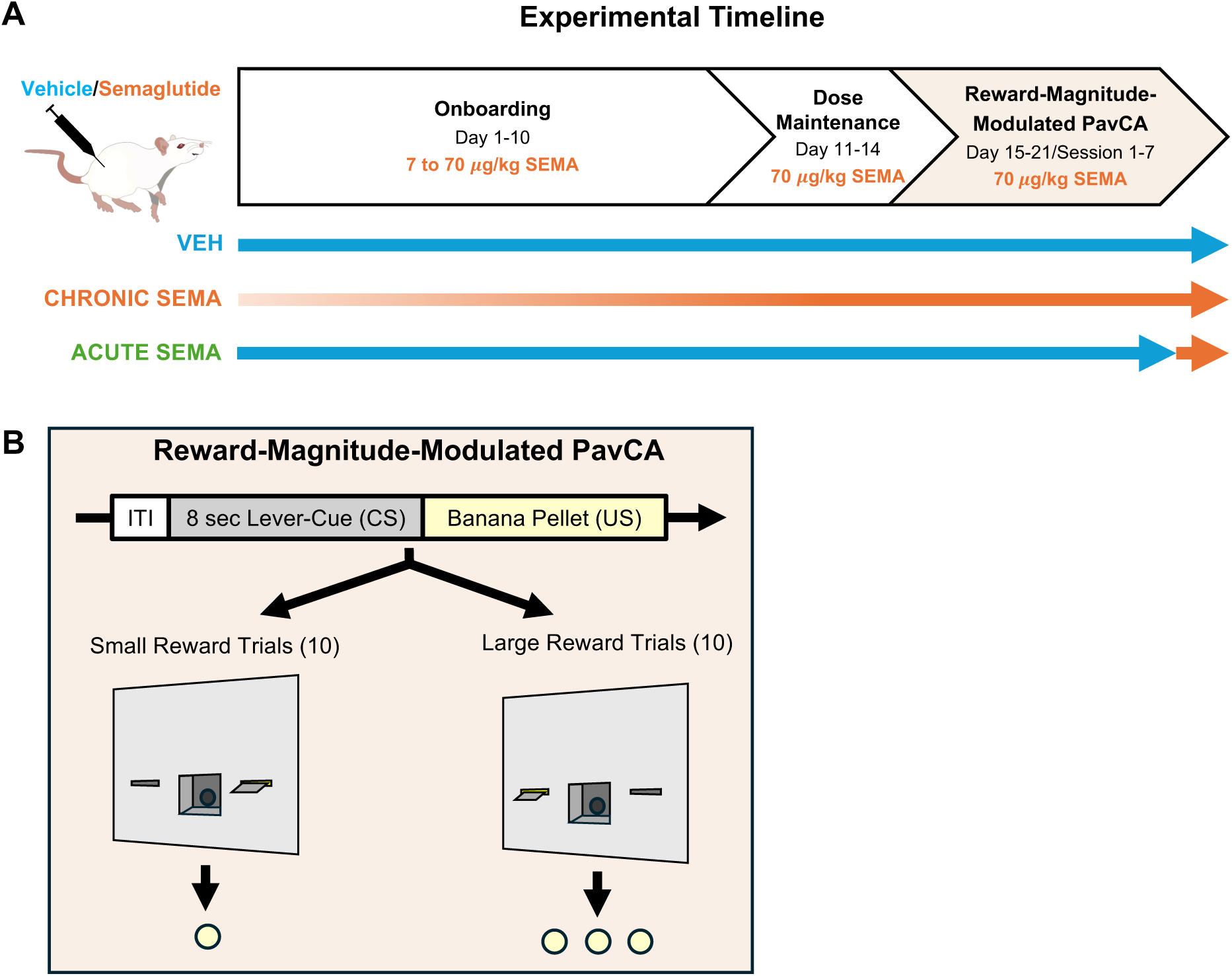
Experimental timeline and reward-magnitude-modulated PavCA (RMM-PavCA) behavioral testing diagram. **(A)** Experimental timeline (days 1-21) across onboarding, dose maintenance, and RMM-PavCA testing phases for vehicle, chronic, and acute semaglutide treatment groups. **(B)** Each RMM-PavCA session consisted of 20 total intermixed trials separated by 30*-*150 sec intertrial intervals (ITIs). For small reward trials (10), 8 sec lever-cue presentation on one side of the food cup predicted a single banana pellet. For large reward trials (10), 8 sec lever-cue presentation on the other side of the food cup predicted 3 banana pellets.

#### Reward pellet exposure

During the exposure phase (experimental days 11-12), approximately 50 45-mg banana-flavored grain pellets (Bio-Serv, Flemington, NJ; Product #F0059) were placed in a cleared corner of each home cage per day to familiarize rats with the food reward, or unconditioned stimulus (US), used during behavioral testing.

#### Reward pellet pretraining

The pretraining phase consisted of two sessions conducted on experimental days 13-14. Rats were placed in behavioral testing chambers, and the house light was illuminated following a 3-min dark period. Twenty-five reward pellets were then delivered according to a variable-interval 30-sec schedule (range 0-60 sec). Each session lasted approximately 12.5 min.

#### Reward-magnitude-modulated Pavlovian conditioned approach (RMM-PavCA) procedures

The RMM-PavCA testing phase included seven daily sessions (sessions 1-7, experimental days 15-21; **Fig. 1A**). Each session began with a 60-sec dark period, after which the house light was illuminated. During small-reward trials (10 per session), a lever-cue on one side of the food cup was inserted and illuminated for 8 sec. Upon lever retraction, one reward pellet was delivered (**Fig. 1B**). During large reward trials (10 per session), the lever-cue on the opposite side of the food cup was inserted and illuminated for 8 sec. Upon lever-cue retraction, three reward pellets were delivered sequentially. Assignment of reward magnitude to the left or right lever-cue was randomized, but remained consistent for each rat. Trials were presented pseudorandomly, with no more than two consecutive trials of the same type, according to a variable-interval 90-sec schedule (range 30-150 sec). After all 20 trials were completed, the houselight was extinguished and the session ended. Each session lasted approximately 33 min.

### Data Analysis and Statistics

#### Change in bodyweight

Change in body weight was calculated relative to baseline body weight prior to treatment initiation (i.e., proportional body-weight change = [(current weight – baseline weight)/baseline weight]).

#### Reward-magnitude-modulated PavCA behavioral measures

Behavioral measures for each RMM-PavCA session included the total number of lever-cue contacts, probability of a lever-cue contact, latency to the first lever-cue contact, total number of food cup entries, probability of a food cup entry, and latency (with maximum of 8 sec) to first food cup entry during lever-cue presentation for each trial type. Probabilities represent the proportion of trials containing at least one response. Latencies were restricted to the 8-sec cue period such that trials without the relevant response were assigned a latency of 8 sec.

A composite PavCA score for each trial type was calculated as the mean of three measures: response bias [(lever-cue contacts – food cup entries)/(lever-cue contacts + food cup entries)], probability difference [(p(lever-cue contact) – p(food cup entry)], and latency score [(average latency to enter the food cup – average latency to lever-cue contact)/duration of the lever-cue presentation] (Meyer et al. 2012). A PavCA score of +1 indicates exclusively sign-tracking behavior, a score of −1 indicates exclusively goal-tracking behavior.

#### Distribution of behaviors across trial types

The proportional distribution of behavior across small and large reward trials during session 7 was determined using lever-cue and food-cup latency data. Each trial was categorized as “Lever-First” if a lever-cue contact preceded a food-cup entry, “Lever-Only” if one or more lever-cue contacts occurred without a food-cup entry, “Cup-First” if a food-cup entry preceded a lever-cue contact, “Cup-Only” if one or more food cup entries occurred without a lever-cue contact, or “None” if neither response occurred (Saunders and Robinson 2012). Trial proportions were averaged within treatment group and reward condition to generate pie charts (**Fig. 4E**). These distributions were examined descriptively and not subjected to statistical analyses.

#### Other behavioral measures

Additionally, the proportion of delivered pellets remaining in the food cup at the end of each session and the rate of food-cup entries during non-cue periods were recorded. For a subset of rats, session 7 was extended from 20 to 40 trials, to determine differential effects with more reward delivery; however, there were no significant effects of this trial extension, so data from trials 21-40 are not included in this study. Pellet consumption and food-cup entry data during the non-cue period from these rats were excluded because the additional trials increased both the number of pellets delivered and the duration of the non-cue period (excluded: VEH n = 5 males and 5 females, CHRONIC SEMA n = 5 males and 6 females, ACUTE SEMA n = 6 males and 5 females). No other measures were affected by this extension, as they were calculated based on the first 20 trials for all rats.

#### Graphing and statistical analyses

Graphs were generated using GraphPad Prism 11.0.2 (GraphPad Software 2026, San Diego, CA). Data were analyzed using SPSS Statistics 31 and 32 (IBM Corp. 2025, 2026, Armonk, NY). Linear mixed models (LMMs), with subject included as a random effect, were used for analyses across multiple time points, and analyses of variance (ANOVAs) were used for outcomes measured at a single time point. The covariance structure for each LMM was selected based on the lowest Akaike information criterion (AIC) value. Statistical significance level was set as p < 0.05, and Bonferroni-corrected post hoc pairwise comparisons were conducted as appropriate.

For body weight analyses, the VEH and future ACUTE SEMA groups were combined into a single vehicle-treated group for experimental days 1-20 because both groups received vehicle during this period. LMM analyses included the fixed effects of sex, experimental day, and experimental group (VEH vs. CHRONIC SEMA), as well as all interactions among these factors. Experimental day was treated as a repeated measure, with subject included as a random effect. For experimental day 21, after administration of acute semaglutide on the day prior, a two-way ANOVA included sex and experimental group (VEH vs. CHRONIC SEMA vs. ACUTE SEMA) as between-subject factors. To directly assess the effects of acute semaglutide treatment on body weight, we compared experimental day 20 and 21 between the VEH and ACUTE SEMA treatment groups. Mixed ANOVAs included sex and experimental group as between-subjects factors and experimental day as within-subjects factors.

For behavioral analyses, the VEH and future ACUTE SEMA groups were combined into a single vehicle-treated group for PavCA sessions 1-6 because both groups received vehicle during this period. LMMs included the fixed effects of sex, session, reward magnitude trial type (large vs. small), and experimental group (VEH vs. CHRONIC SEMA), as well as all interactions among these factors. Session and reward magnitude were treated as repeated measures, and subject was included as a random effect. For PavCA session 7, after administration of acute semaglutide the day prior, mixed ANOVAs included sex and experimental group (VEH vs.

CHRONIC SEMA vs. ACUTE SEMA) as between-subjects factors and reward magnitude trial type (large vs. small) as a within-subjects factor. Additionally, to directly assess the effects of acute semaglutide treatment, we compared behavior across PavCA session 6 and 7 between the VEH and ACUTE SEMA treatment groups. Mixed ANOVAs included sex and experimental group as between-subjects factors and reward magnitude trial type (large vs. small) and session as within-subjects factors. Reward magnitude trial type was not included as a factor in analyses for the proportion of pellets remaining or the rate of non-cue food-cup entries.

## Results

### Change in body weight

Chronic semaglutide reduced the proportional change in body weight relative to vehicle treatment across dose-escalation and maintenance days 1-20 (main effect of treatment: F_1,67.531_ = 267.985, p < 0.001; **Fig. 2A**; see **Table 1** for full statistical results). Specifically, chronic treatment resulted in weight loss in females (sex x treatment interaction: F_1,67.531_ = 17.518, p < 0.001; post-hoc treatment comparison: p < 0.001) and attenuated weight gain in males (post-hoc treatment comparison: p < 0.001).

**Fig. 2.**
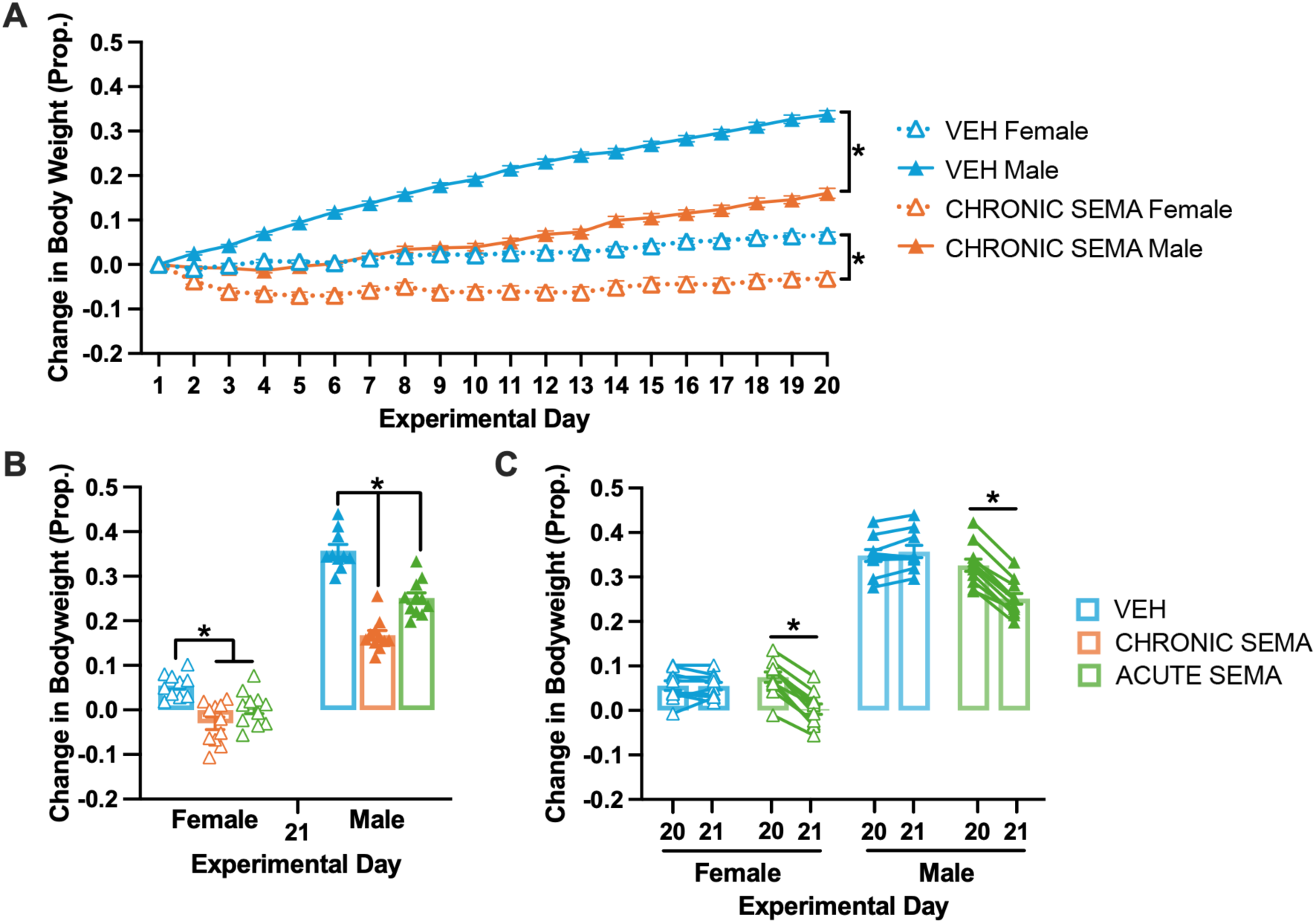
Both chronic and acute semaglutide treatment decrease change in body weight from baseline. **(A)** Line graph shows change in body weight as a proportion of baseline (mean +/- SEM) across experimental days 1*-*20, comparing chronic semaglutide and vehicle treatment in male and female subjects (*indicates p < 0.05 for post-hoc treatment comparison within sex). **(B)** Bar graph shows the same change in body weight measure (mean +/- SEM; triangular points represent individual subjects) on experimental day 21, including the acute semaglutide treatment group (*indicates p < 0.05 for post-hoc treatment comparison within sex). **(C)** Bar graph shows change in body weight measure (mean +/- SEM; triangular points represent individual subjects) over experimental days 20-21, including the vehicle and acute semaglutide treatment groups (*indicates p < 0.05 for post-hoc session comparison within treatment group).

**Table 1.** LMM (experimental days 1-20) and ANOVA (experimental day 21; experimental day 20-21) statistical results for change in body weight data shown in Fig 2. Significant effects (p < 0.05) are bolded and italicized.

|  | Change in Body weight<br>Days 1-20 | Change in Body weight<br>Day 21 | Change in Body weight<br>Day 20-21 |
| --- | --- | --- | --- |
| Sex | ***$F(1,67.531)=$***<br>***$460.302, p<0.001$*** | ***$F(1,58)=$***<br>***$659.037, p<0.001$*** | ***$F(1,38)=$***<br>***$536.446, p<0.001$*** |
| Treatment | ***$F(1,67.531)=$***<br>***$267.985, p<0.001$*** | ***$F(2,58)=$***<br>***$65.809, p<0.001$*** | ***$F(1,38)=$***<br>***$11.558, p=0.002$*** |
| Session | ***$F(19,162.498)=$***<br>***$77.092, p<0.001$*** | NA | ***$F(1,38)=$***<br>***$176.669, p<0.001$*** |
| Sex*<br>Treatment | ***$F(1,67.531)=$***<br>***$17.518, p<0.001$*** | ***$F(2,58)=$***<br>***$9.594, p<0.001$*** | $F(1,38)=$<br>4.069, $p=0.051$ |
| Sex*Session | ***$F(19,162.498)=$***<br>***$51.393, p<0.001$*** | NA | $F(1,38)=$<br>0.391, $p=0.535$ |
| Treatment*<br>Session | ***$F(19,162.498)=$***<br>***$22.036, p<0.001$*** | NA | ***$F(1,38)=$***<br>***$220.872, p<0.001$*** |
| Sex*<br>Treatment*<br>Session | ***$F(19,162.498)=$***<br>***$3.641, p<0.001$*** | NA | $F(1,38)=$<br>1.726, $p=0.197$ |

Following PavCA session 6 (experimental session 20), vehicle-treated rats were divided into VEH and ACUTE SEMA groups. Rats in the ACUTE SEMA group, received a single dose of semaglutide approximately 24 h before body-weight measurement and behavioral testing on experimental day 21. The pattern observed across days 1-20 continued on day 21, with chronic semaglutide producing a greater reduction in proportional body weight relative to vehicle treatment (main effect of treatment: F_2,58_ = 65.809, p < 0.001; sex x treatment interaction: F_2,58_ = 9.594, p < 0.001; post-hoc pairwise comparisons: CHRONIC SEMA vs. VEH, p < 0.001 for both males and females; **Fig. 2B**; **Table 1**). Acute semaglutide treatment also resulted in lower change in body weight relative to vehicle treatment in both females (post-hoc pairwise comparison: ACUTE SEMA vs. VEH, p = 0.010) and males (ACUTE SEMA vs. VEH, p < 0.001). Change in body weight after acute treatment was not significantly different from chronic treatment in females (post-hoc pairwise comparison: ACUTE SEMA vs. CHRONIC SEMA, p = 0.151). In males, the change in bodyweight after acute treatment was significantly different from chronic treatment, with the proportional change higher in the ACUTE SEMA group relative to CHRONIC SEMA (ACUTE SEMA vs. CHRONIC SEMA, p < 0.001; **Fig 2B**). Comparing across experimental days 20-21, ACUTE SEMA reduced body weight within subjects, whereas there was not a significant change in the proportional body weight of VEH-treated rats (treatment x session interaction: F_1,38_ = 220.872, p < 0.001; post-hoc pairwise comparisons for days 20-21: ACUTE SEMA, p < 0.001; VEH, p = 0.285; **Fig. 2C**, **Table 1**). Thus, both chronic and acute semaglutide treatment significantly reduced body weight relative to vehicle treatment.

### Chronic semaglutide effects on reward-magnitude-modulated PavCA behavior

Beginning on experimental day 15, all rats underwent RMM-PavCA testing. Presentation of a lever-cue on one side of the food cup predicted a small reward of one banana-flavored pellet, whereas presentation of a lever-cue on the opposite side predicted a large reward of three banana-flavored pellets (**Fig. 1B**). Across RMM-PavCA sessions 1-6, rats exhibited more sign-tracking behaviors during large-than small-reward trials, including more lever-cue contacts (main effect of reward magnitude: F_1,57.951_ = 6.321, p = 0.015; **Fig. 3A**; **Table 2** for full statistical results), a greater probability of lever-cue contact (F_1,61.463_ = 6.359, p = 0.014; **Fig. 3B**), and a shorter latency to contact the lever-cue (F_1,60.754_ = 12.564, p < 0.001; **Fig 3C**). Chronic semaglutide treatment also increased sign-tracking overall, as indicated by more lever-cue contacts (main effect of treatment: F_1,60.001_ = 13.391, p < 0.001), a greater probability of lever-cue contact (main effect of treatment: F_1,66.734_ = 15.194, p < 0.001), and a shorter latency to contact the lever-cue (main effect of treatment: F_1,65.880_ = 22.762, p < 0.001). There was a significant sex x session x treatment interaction for probability of lever-cue contact (F_5,139.853_ = 2.854, p=0.017), with VEH-treated females having higher probability of contact than VEH-treated males on sessions 2-4 (post-hoc comparisons of sex within treatment: sessions 2-4, p < 0.03). There were no other significant effects of sex on lever-directed behaviors across RMM-PavCA sessions 1-6.

**Fig. 3.**
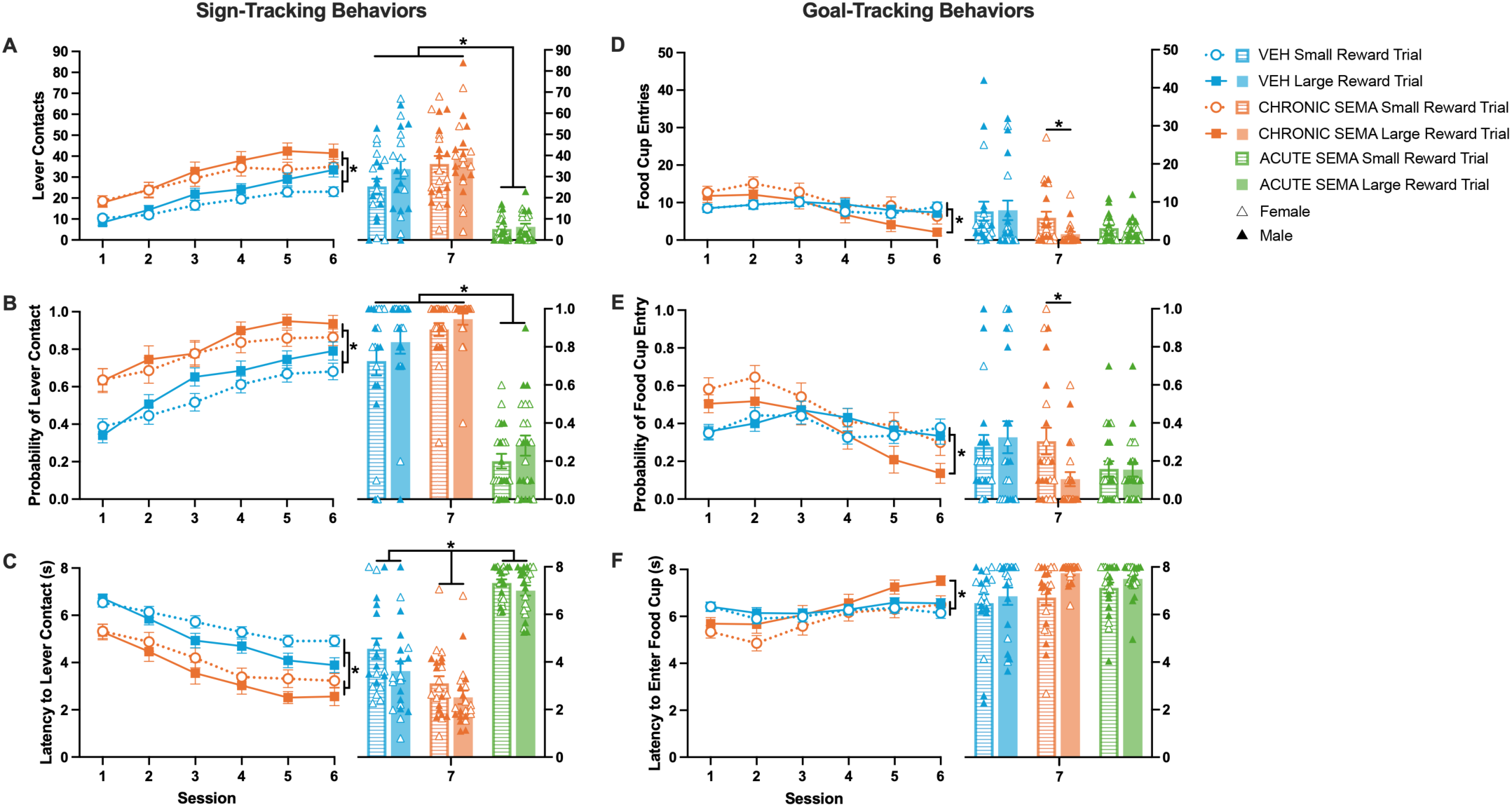
Chronic and acute semaglutide treatment have distinct effects on lever-cue directed behavior. Line graphs show behavioral data (mean +/- SEM) across sessions 1*-*6 of reward-magnitude-modulated PavCA (RMM-PavCA), comparing chronic semaglutide and vehicle treatment for small and large reward trials. Bar graphs show the same behavioral measures (mean +/- SEM; triangular points represent individual subjects) on session 7 of RMM-PavCA, including the acute semaglutide treatment group. Lever-cue directed (i.e. sign-tracking) behavioral measures include **(A)** contacts, **(B)** probability of contact, and **(C)** latency to contact the lever-cue (for A*-*C line graphs: *indicates p < 0.05 for main effect of treatment; for bar graphs: *indicates p < 0.05 for post-hoc treatment comparison). Food cup directed (i.e. goal-tracking) behaviors included **(D)** entries, **(E)** probability of entry, and **(F)** latency to enter the food cup (for D*-*F line and bar graphs: *indicates p < 0.05 for post-hoc reward magnitude comparison within treatment group).

**Table 2.**
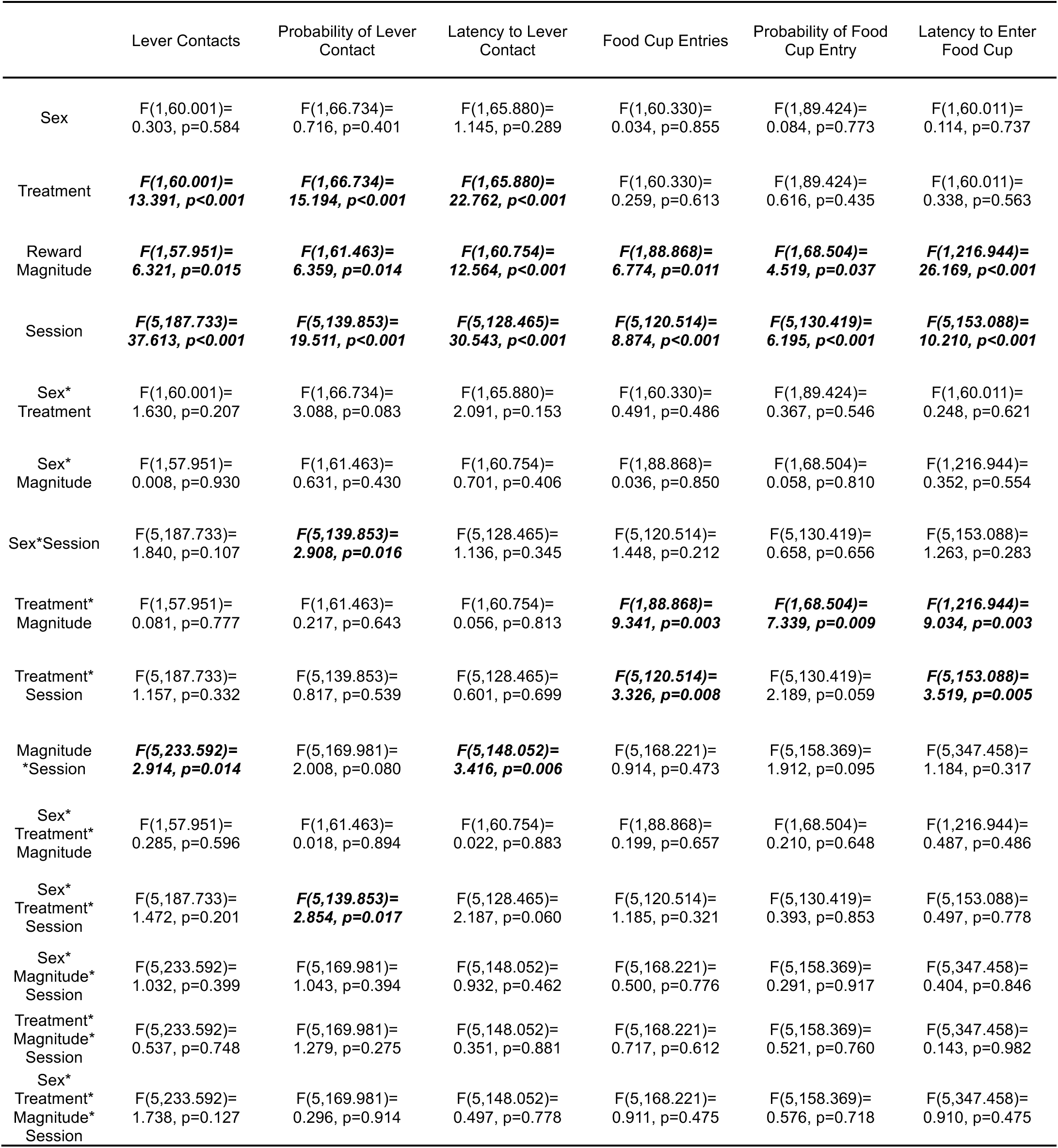
LMM statistical results for sessions 1-6 reward-magnitude-modulated PavCA behavioral data shown in Fig. 3. Significant effects (p < 0.05) are bolded and italicized.

|  | Lever Contacts | Probability of Lever Contact | Latency to Lever Contact | Food Cup Entries | Probability of Food Cup Entry | Latency to Enter Food Cup |
| --- | --- | --- | --- | --- | --- | --- |
| Sex | $F(1,60.001)=0.303, p=0.584$ | $F(1,66.734)=0.716, p=0.401$ | $F(1,65.880)=1.145, p=0.289$ | $F(1,60.330)=0.034, p=0.855$ | $F(1,89.424)=0.084, p=0.773$ | $F(1,60.011)=0.114, p=0.737$ |
| Treatment | ***$F(1,60.001)=13.391, p<0.001$*** | ***$F(1,66.734)=15.194, p<0.001$*** | ***$F(1,65.880)=22.762, p<0.001$*** | $F(1,60.330)=0.259, p=0.613$ | $F(1,89.424)=0.616, p=0.435$ | $F(1,60.011)=0.338, p=0.563$ |
| Reward Magnitude | ***$F(1,57.951)=6.321, p=0.015$*** | ***$F(1,61.463)=6.359, p=0.014$*** | ***$F(1,60.754)=12.564, p<0.001$*** | ***$F(1,88.868)=6.774, p=0.011$*** | ***$F(1,68.504)=4.519, p=0.037$*** | ***$F(1,216.944)=26.169, p<0.001$*** |
| Session | ***$F(5,187.733)=37.613, p<0.001$*** | ***$F(5,139.853)=19.511, p<0.001$*** | ***$F(5,128.465)=30.543, p<0.001$*** | ***$F(5,120.514)=8.874, p<0.001$*** | ***$F(5,130.419)=6.195, p<0.001$*** | ***$F(5,153.088)=10.210, p<0.001$*** |
| Sex* Treatment | $F(1,60.001)=1.630, p=0.207$ | $F(1,66.734)=3.088, p=0.083$ | $F(1,65.880)=2.091, p=0.153$ | $F(1,60.330)=0.491, p=0.486$ | $F(1,89.424)=0.367, p=0.546$ | $F(1,60.011)=0.248, p=0.621$ |
| Sex* Magnitude | $F(1,57.951)=0.008, p=0.930$ | $F(1,61.463)=0.631, p=0.430$ | $F(1,60.754)=0.701, p=0.406$ | $F(1,88.868)=0.036, p=0.850$ | $F(1,68.504)=0.058, p=0.810$ | $F(1,216.944)=0.352, p=0.554$ |
| Sex*Session | $F(5,187.733)=1.840, p=0.107$ | ***$F(5,139.853)=2.908, p=0.016$*** | $F(5,128.465)=1.136, p=0.345$ | $F(5,120.514)=1.448, p=0.212$ | $F(5,130.419)=0.658, p=0.656$ | $F(5,153.088)=1.263, p=0.283$ |
| Treatment* Magnitude | $F(1,57.951)=0.081, p=0.777$ | $F(1,61.463)=0.217, p=0.643$ | $F(1,60.754)=0.056, p=0.813$ | ***$F(1,88.868)=9.341, p=0.003$*** | ***$F(1,68.504)=7.339, p=0.009$*** | ***$F(1,216.944)=9.034, p=0.003$*** |
| Treatment* Session | $F(5,187.733)=1.157, p=0.332$ | $F(5,139.853)=0.817, p=0.539$ | $F(5,128.465)=0.601, p=0.699$ | ***$F(5,120.514)=3.326, p=0.008$*** | $F(5,130.419)=2.189, p=0.059$ | ***$F(5,153.088)=3.519, p=0.005$*** |
| Magnitude*Session | ***$F(5,233.592)=2.914, p=0.014$*** | $F(5,169.981)=2.008, p=0.080$ | ***$F(5,148.052)=3.416, p=0.006$*** | $F(5,168.221)=0.914, p=0.473$ | $F(5,158.369)=1.912, p=0.095$ | $F(5,347.458)=1.184, p=0.317$ |
| Sex* Treatment* Magnitude | $F(1,57.951)=0.285, p=0.596$ | $F(1,61.463)=0.018, p=0.894$ | $F(1,60.754)=0.022, p=0.883$ | $F(1,88.868)=0.199, p=0.657$ | $F(1,68.504)=0.210, p=0.648$ | $F(1,216.944)=0.487, p=0.486$ |
| Sex* Treatment* Session | $F(5,187.733)=1.472, p=0.201$ | ***$F(5,139.853)=2.854, p=0.017$*** | $F(5,128.465)=2.187, p=0.060$ | $F(5,120.514)=1.185, p=0.321$ | $F(5,130.419)=0.393, p=0.853$ | $F(5,153.088)=0.497, p=0.778$ |
| Sex* Magnitude* Session | $F(5,233.592)=1.032, p=0.399$ | $F(5,169.981)=1.043, p=0.394$ | $F(5,148.052)=0.932, p=0.462$ | $F(5,168.221)=0.500, p=0.776$ | $F(5,158.369)=0.291, p=0.917$ | $F(5,347.458)=0.404, p=0.846$ |
| Treatment* Magnitude* Session | $F(5,233.592)=0.537, p=0.748$ | $F(5,169.981)=1.279, p=0.275$ | $F(5,148.052)=0.351, p=0.881$ | $F(5,168.221)=0.717, p=0.612$ | $F(5,158.369)=0.521, p=0.760$ | $F(5,347.458)=0.143, p=0.982$ |
| Sex* Treatment* Magnitude* Session | $F(5,233.592)=1.738, p=0.127$ | $F(5,169.981)=0.296, p=0.914$ | $F(5,148.052)=0.497, p=0.778$ | $F(5,168.221)=0.911, p=0.475$ | $F(5,158.369)=0.576, p=0.718$ | $F(5,347.458)=0.910, p=0.475$ |

For goal-tracking measures, the effect of chronic semaglutide differed as a function of reward magnitude across sessions 1-6. Relative to small-reward trials, rats treated chronically with semaglutide exhibited fewer food cup entries during large-reward trials (treatment x reward magnitude interaction: F_1,88.868_ = 9.341, p = 0.003; post-hoc pairwise comparisons for small vs. large trial type: CHRONIC SEMA, p < 0.001; VEH, p = 0.700; **Fig. 3D**), a lower probability of food cup entry (treatment x reward magnitude interaction: F_1,68.504_ = 7.339, p = 0.009; post-hoc pairwise comparisons for small vs. large trial type: CHRONIC SEMA, p = 0.004; VEH, p = 0.621; **Fig. 3E**), and a longer latency to enter the food cup (treatment x reward magnitude interaction: F_1,216.944_ = 9.034, p = 0.003; post-hoc pairwise comparisons for small vs. large trial type: CHRONIC SEMA, p < 0.001; VEH, p = 0.073; **Fig. 3F**). This pattern is consistent with greater allocation of behavior to the lever-cue rather than the food-cup during large-reward trials in the chronic semaglutide treatment group. There were no significant effects of sex on any food cup-directed behaviors across sessions 1-6.

### Acute semaglutide effects on reward-magnitude-modulated PavCA behavior

For RMM-PavCA session 7, rats previously treated with vehicle were divided into VEH and ACUTE SEMA groups, with the latter receiving a single dose of semaglutide approximately 24 h before the final session. In contrast to the effects observed following chronic treatment, rats receiving acute semaglutide exhibited lower levels of lever-cue directed behavior, including fewer lever-cue contacts (main effect of treatment: F_2,58_ = 38.288, p < 0.001; post-hoc pairwise comparisons: ACUTE SEMA vs. VEH, p < 0.001; ACUTE SEMA vs. CHRONIC SEMA, p < 0.001; **Fig. 3A**; see **Table 3** for full statistical results), a lower probability of lever-cue contact (main effect of treatment: F_2,58_ = 70.621, p < 0.001; post-hoc pairwise comparisons: ACUTE SEMA vs. VEH, p < 0.001; ACUTE SEMA vs. CHRONIC SEMA, p < 0.001; **Fig. 3B**), and a longer latency to contact the lever-cue (main effect of treatment: F_2,58_ = 85.579, p < 0.001; post-hoc pairwise comparisons: ACUTE SEMA vs. VEH, p < 0.001; ACUTE SEMA vs. CHRONIC SEMA, p < 0.001; VEH vs. CHRONIC SEMA, p = 0.002; **Fig. 3C**). Acute semaglutide treatment did not significantly affect goal-tracking measures on session 7 (all main effects of treatment: p > 0.05; **Fig. 3D-F**). There were no significant effects of sex for any measures on session 7.

**Table 3.** ANOVA statistical results for session 7 reward-magnitude-modulated PavCA behavioral data shown in Fig. 3. Significant effects (p < 0.05) are bolded and italicized.

|  | Lever Contacts | Probability of Lever Contact | Latency to Lever Contact | Food Cup Entries | Probability of Food Cup Entry | Latency to Enter Food Cup |
| --- | --- | --- | --- | --- | --- | --- |
| Sex | F(1,58)=<br>0.003, $p=0.954$ | F(1,58)=<br>0.007, $p=0.932$ | F(1,58)=<br>0.750, $p=0.390$ | F(1,58)=<br>1.076, $p=0.304$ | F(1,58)=<br>1.231, $p=0.272$ | F(1,58)=<br>1.525, $p=0.222$ |
| Treatment | ***F(2,58)=***<br>***38.288, $p<0.001$*** | ***F(2,58)=***<br>***70.621, $p<0.001$*** | ***F(2,58)=***<br>***85.579, $p<0.001$*** | F(2,58)=<br>2.968, $p=0.059$ | F(2,58)=<br>2.122, $p=0.129$ | F(2,58)=<br>2.514, $p=0.090$ |
| Reward Magnitude | F(1,58)=<br>3.261, $p=0.076$ | ***F(1,58)=***<br>***6.770, $p=0.012$*** | ***F(1,58)=***<br>***8.525, $p=0.005$*** | ***F(1,58)=***<br>***5.137, $p=0.027$*** | F(1,58)=<br>2.292, $p=0.135$ | ***F(1,58)=***<br>***13.743, $p<0.001$*** |
| Sex*<br>Treatment | F(2,58)=<br>0.944, $p=0.395$ | F(2,58)=<br>0.958, $p=0.390$ | F(2,58)=<br>2.035, $p=0.140$ | F(2,58)=<br>0.940, $p=0.397$ | F(2,58)=<br>0.560, $p=0.574$ | F(2,58)=<br>1.130, $p=0.330$ |
| Sex*<br>Magnitude | F(1,58)=<br>0.052, $p=0.820$ | F(1,58)=<br>0.137, $p=0.713$ | F(1,58)=<br>0.005, $p=0.943$ | F(1,58)=<br>0.022, $p=0.883$ | F(1,58)=<br>0.103, $p=0.750$ | F(1,58)=<br>0.006, $p=0.939$ |
| Treatment*<br>Magnitude | F(2,58)=<br>0.872, $p=0.424$ | F(2,58)=<br>0.189, $p=0.829$ | F(2,58)=<br>0.712, $p=0.495$ | ***F(2,58)=***<br>***3.896, $p=0.026$*** | ***F(2,58)=***<br>***4.996, $p=0.010$*** | F(2,58)=<br>2.367, $p=0.103$ |
| Sex*<br>Treatment*<br>Magnitude | F(2,58)=<br>0.538, $p=0.587$ | F(2,58)=<br>0.518, $p=0.598$ | F(2,58)=<br>0.016, $p=0.984$ | F(2,58)=<br>0.058, $p=0.943$ | F(2,58)=<br>0.014, $p=0.986$ | F(2,58)=<br>0.015, $p=0.985$ |

In agreement with the data above, when we compared across sessions 6-7 within subjects, ACUTE SEMA reduced all lever-directed behaviors, including lower contacts (treatment x session interaction: F_1,38_ = 53.349, p < 0.001; post-hoc pairwise comparisons for sessions 6-7: ACUTE SEMA, p < 0.001; VEH, p = 0.268; **Fig. S1A**; **Table S1** for full statistical results), lower probability (treatment x session interaction: F_1,38_ = 115.632, p < 0.001; post-hoc pairwise comparisons for sessions 6-7: ACUTE SEMA, p < 0.001; VEH, p = 0.393; **Fig. S1B**), and increased latency (treatment x session interaction: F_1,38_ = 97.788, p < 0.001; post-hoc pairwise comparisons for sessions 6-7: ACUTE SEMA, p < 0.001; VEH, p = 0.121; **Fig. S1C**). ACUTE SEMA also reduced food cup entries (treatment x session interaction: F_1,38_ = 7.195, p = 0.011; post-hoc pairwise comparisons for sessions 6-7: ACUTE SEMA, p < 0.001; VEH, p = 0.311; **Fig. S1D**), with no significant effects on probability or latency to enter (treatment x session interactions: p > 0.130; **Fig. S1E-F**) and no significant sex effects across these measures.

Composite measures from session 7 further characterized the relative difference in sign-versus goal-tracking tendencies (**Fig. 4**). Acute semaglutide resulted in lower scores reflecting less sign-tracking relative to goal-tracking, as indicated by lower response bias (main effect of treatment: F_2,58_ = 8.528, p < 0.001; post-hoc pairwise comparisons: ACUTE SEMA vs.

**Fig. 4.**
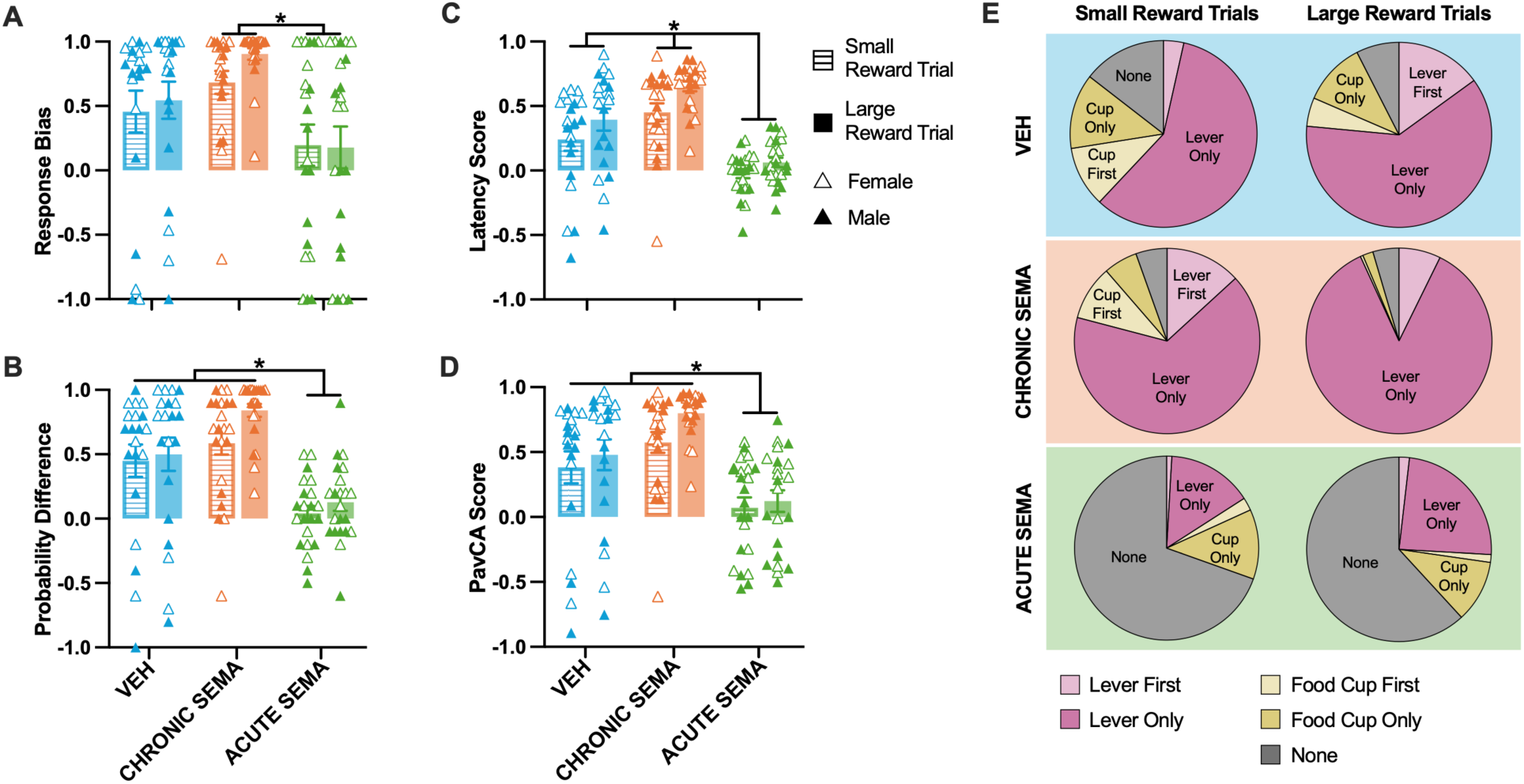
Acute semaglutide treatment decreases sign-tracking relative to goal-tracking behaviors. Bar graphs show behavioral data (mean +/- SEM; triangular points represent individual subjects) on session 7 of reward-magnitude-modulated PavCA (RMM-PavCA), comparing chronic semaglutide, acute semaglutide, and vehicle treatment for small and large reward trials. For all behaviors including **(A)** response bias, **(B)** probability difference, **(C)** latency difference, and **(D)** overall PavCA score (an average of A*-*C), +1 indicates exclusively sign-tracking responses and −1 indicates exclusively goal-tracking responses. *Indicates p < 0.05 for post-hoc treatment comparison. **(E)** Pie charts show the mean distribution of behavior for each treatment group across small and large reward trials, including trials in which subjects interact with the lever-cue first or only the lever-cue (i.e. sign-tracking), the food cup first or only the food cup (i.e. goal-tracking), or did not interact with either the lever-cue or food cup (i.e. none).

CHRONIC SEMA, p < 0.001; **Fig. 4A**; **Table 4** for full statistical results), probability difference (main effect of treatment: F_2,58_ = 15.292, p < 0.001; post-hoc pairwise comparisons: ACUTE SEMA vs. VEH, p = 0.004; ACUTE SEMA vs. CHRONIC SEMA, p < 0.001; **Fig. 4B**), latency score (main effect of treatment: F_2,58_ = 27.409, p < 0.001; post-hoc pairwise comparisons: ACUTE SEMA vs. VEH, p < 0.001; ACUTE SEMA vs. CHRONIC SEMA, p < 0.001; VEH vs.

**Table 4.** ANOVA statistical results for session 7 reward-magnitude-modulated PavCA composite measures shown in Fig. 4A-D. Significant effects (p < 0.05) are bolded and italicized.

|  | Response Bias | Probability Difference | Latency Score | PavCA Score |
| --- | --- | --- | --- | --- |
| Sex | $F(1,58)=$<br>1.180, $p=0.282$ | $F(1,58)=$<br>0.618, $p=0.435$ | $F(1,58)=$<br>1.405, $p=0.241$ | $F(1,58)=$<br>1.053, $p=0.309$ |
| Treatment | ***$F(2,58)=$***<br>***8.528, $p<0.001$*** | ***$F(2,58)=$***<br>***15.292, $p<0.001$*** | ***$F(2,58)=$***<br>***27.409, $p<0.001$*** | ***$F(2,58)=$***<br>***15.429, $p<0.001$*** |
| Reward<br>Magnitude | $F(1,58)=$<br>1.024, $p=0.316$ | ***$F(1,58)=$***<br>***7.494, $p=0.008$*** | ***$F(1,58)=$***<br>***17.741, $p<0.001$*** | ***$F(1,58)=$***<br>***5.088, $p=0.028$*** |
| Sex*<br>Treatment | $F(2,58)=$<br>1.267, $p=0.289$ | $F(2,58)=$<br>0.707, $p=0.498$ | $F(2,58)=$<br>2.100, $p=0.132$ | $F(2,58)=$<br>1.051, $p=0.356$ |
| Sex*<br>Magnitude | $F(1,58)=$<br>0.153, $p=0.697$ | $F(1,58)=$<br>0.269, $p=0.606$ | $F(1,58)=$<br>0.077, $p=0.783$ | $F(1,58)=$<br>0.008, $p=0.929$ |
| Treatment*<br>Magnitude | $F(2,58)=$<br>0.534, $p=0.589$ | $F(2,58)=$<br>1.826, $p=0.170$ | $F(2,58)=$<br>1.276, $p=0.287$ | $F(2,58)=$<br>0.908, $p=0.409$ |
| Sex*<br>Treatment*<br>Magnitude | $F(2,58)=$<br>0.082, $p=0.921$ | $F(2,58)=$<br>0.212, $p=0.810$ | $F(2,58)=$<br>0.097, $p=0.908$ | $F(2,58)=$<br>0.056, $p=0.945$ |

CHRONIC SEMA, p = 0.008; **Fig. 4C**), and overall PavCA score (main effect of treatment: F_2,58_ = 15.429, p < 0.001; ACUTE SEMA vs. VEH, p = 0.010; ACUTE SEMA vs. CHRONIC SEMA, p < 0.001; **Fig. 4D**). Because acute semaglutide did not increase goal-tracking (**Fig. 3**), these lower composite scores primarily reflect lower sign-tracking, rather than a shift toward greater goal-tracking.

Across treatment groups, rats exhibited greater sign-tracking during large- than small-reward trials, as indicated by a greater probability difference (main effect of reward magnitude: F_1,58_ = 7.494, p = 0.008), latency score (main effect of reward magnitude: F_1,58_ = 17.741, p < 0.001), and composite PavCA score on RMM-PavCA session 7 (main effect of reward magnitude: F_1,58_ = 5.088, p = 0.028). There were no significant effects of sex on any behavioral measures on session 7.

### Distribution of behaviors across trial types

To further characterize group differences during RMM-PavCA session 7, we descriptively examined the distribution of behavior across small- and large-reward trial types (**Fig. 4E**). All treatment groups exhibited a greater proportion of lever-directed trials for cues predicting the large reward than for cues predicting the small reward. In the VEH group, among trials in which both lever-cue contacts and food-cup entries occurred, rats were more likely to enter the food cup before contacting the lever-cue during small-reward trials (11% Cup-First vs. 4% Lever First). In contrast, they were more likely to contact the lever-cue first during large-reward trials (5% Cup First vs. 15% Lever First). In the CHRONIC SEMA group, the proportion of trials involving only lever-cue contacts was greater during large-reward trials (86% Lever Only) than small-reward trials (66% Lever Only).

Notably, rats in the ACUTE SEMA group did not contact either the lever-cue or the food cup during most small-reward (70% None) and large-reward (62% None) trials. Despite this generally low level of responding, the proportion of Lever-Only trials was greater for large-reward cues (24%) than for small-reward cues (15%) in the ACUTE SEMA group.

These descriptive data were consistent with greater sign-tracking during large- than small-reward trials across groups. Chronic semaglutide was associated with greater lever-cue directed behaviors overall, whereas acute semaglutide was associated with substantially lower levels of responding.

### Other behavioral measures

To further characterize the effects of semaglutide treatment beyond cue-elicited responding, we examined the proportion of delivered reward pellets remaining in the food cup at the end of each session and the rate of food-cup entries during the non-CS intertrial-interval (ITI) periods. There were no significant effects of sex on pellet consumption or the rate of food-cup entries. The CHRONIC SEMA and VEH groups did not differ in the proportion of pellets remaining across sessions 1-6 of RMM-PavCA testing (main effect of treatment: F_1,61.067_ = 0.957, p = 0.332; **Fig. S2B**; **Table S2** for full statistical results). The groups also did not differ significantly in pellet consumption during the earliest phases of training, averaged across pretraining or on session 1 (p > 0.05; **Fig.S2A**). During PavCA session 7, however, the ACUTE SEMA group left a greater proportion of delivered pellets uneaten than did the CHRONIC SEMA and VEH groups (main effect of treatment: F_2,26_ = 13.812, p < 0.001; post-hoc pairwise comparisons: ACUTE SEMA vs. VEH, p < 0.001; ACUTE SEMA vs. CHRONIC SEMA, p < 0.001; **Fig. S2B**). No rats in the VEH or CHRONIC SEMA groups left pellets uneaten during this session, while 9 of 11 rats (3 of 5 males, 6 of 6 females) in the ACUTE SEMA group did not consume all of the delivered pellets.

The rate of entry into the food cup during non-CS periods differed between the CHRONIC SEMA and VEH groups only during the earliest phase of PavCA training (treatment x session interaction: F_5,99.932_ = 3.905, p = 0.003). Specifically, the CHRONIC SEMA group exhibited a higher non-CS food-cup entry rate than the VEH group during session 1 (p = 0.001), whereas the groups did not differ during subsequent sessions (all p > 0.255; **Fig. S2D**). Across pretraining and session 1, chronic semaglutide was similarly associated with a higher non-CS food-cup entry rate (main effect of treatment: F_1,60_ = 28.235, p < 0.001; **Fig. S2C**). This group difference was no longer evident during subsequent sessions, presumably after the cue-reward relationship was learned.

During session 7, the ACUTE SEMA group exhibited a lower non-CS food-cup entry rate than the CHRONIC SEMA group (main effect of treatment: F_2,26_ = 3.631, p = 0.041; post-hoc pairwise comparison: ACUTE SEMA vs. CHRONIC SEMA p = 0.040; **Fig. S2D**). The ACUTE SEMA and VEH groups did not differ significantly. Together with the high proportion of trials without a lever-cue contact or food-cup entry (**Fig. 4E**), this finding is consistent with a broader reduction in behavioral responding following acute treatment, that was not restricted to the reward-predictive cue.

## Discussion

Although both treatment regimens reduced body weight relative to vehicle treatment, acute and chronic semaglutide regimens produced distinct behavioral outcomes in a reward-magnitude-modulated PavCA paradigm. As expected, rats generally exhibited greater sign-tracking toward cues that predict larger rewards. This finding suggests that sign-tracking is a behavioral representation of motivational features of the outcome being attributed to the cue, and is largely consistent with the incentive salience theory (Berridge 2023). Chronic semaglutide treatment increased overall sign-tracking measures while decreasing goal-tracking during trials involving the large-reward-predictive lever-cue. This pattern is consistent with greater allocation of behavior toward the lever-cue and away from the food cup. However, because semaglutide did not differentially alter the effect of reward magnitude on sign-tracking, the findings indicate an overall increase in cue-directed behavior rather than selective enhancement of the large-reward cue.

This effect may reflect increased motivational value of the reward itself. Consistent with this possibility, chronic semaglutide treatment increased food cup responding during pretraining and early PavCA sessions, before the cue-reward association was fully established. Once this association was formed during later sessions, responding was increasingly directed toward the predictive lever-cue. This temporal pattern is consistent with enhanced food-cup-directed responding that subsequently transferred to the reward-predictive cue, although general activity and reward motivation were not independently measured. This interpretation is further supported by prior findings from our laboratory demonstrating that chronic semaglutide treatment increased conditioned reinforcement for reward-associated cues as well as progressive ratio responding for the same palatable food reward (Chang et al. 2026). However, we did not previously observe changes in sign- or goal-tracking behavior during standard PavCA training following chronic semaglutide treatment (Chang et al. 2026). The present within-subjects RMM-PavCA design, which included cues predicting different reward magnitudes, seems to have revealed treatment-related differences that were not captured by standard PavCA procedures.

In contrast to chronic treatment, acute administration of semaglutide prior to the final reward-magnitude-modulated -PavCA session was associated with a decrease in all sign-tracking measures, and a greater proportion of trials during which rats made no response. Acute treatment was also associated with lower pellet consumption and food cup responding during the intertrial interval. Together, these measures indicate a broader reduction in food- and cue-directed behavior that was not restricted to sign-tracking. This pattern may reflect malaise or other adverse effects associated with initial semaglutide exposure, although this requires further investigation as malaise and locomotor activity were not directly measured in the current study.

Semaglutide treatment in humans is known to cause adverse gastrointestinal effects, with one study reporting nausea in 43.9% of participants, most commonly during the early dose-escalation phase (Wharton et al. 2022a). Related effects have been reported in preclinical models. Short-term semaglutide treatment in rats increased kaolin clay consumption, a measure of visceral malaise in rodents, while decreasing operant responding for sucrose, and increasing aversive orofacial responses to a novel palatable taste (Ghidewon et al. 2022). Additionally, in mice, a single dose of semaglutide administered after sucrose exposure led to conditioned taste aversion as measured by reduced sucrose preference 2 days after conditioning despite no further drug exposure (Teixidor-Deulofeu et al. 2025). It is possible, therefore, that gastrointestinal or aversive effects may have contributed to the lower food-cue-directed behavior observed after acute semaglutide treatment. However, the present measures cannot distinguish these effects from reduced appetite, altered incentive motivation, or other forms of behavioral suppression.

Together, our findings and other preclinical evidence raise the possibility that adverse effects contribute to changes in food-cue reactivity following acute or initial GLP-1RA administration. These findings may be relevant to reports of decreased “food-noise” in individuals taking GLP-1RAs (de Vere Hunt et al. 2026; Rutigliani and Mattes 2026). In humans, one study found that liraglutide treatment in individuals with diabetes reduced neural responses to images of highly desired foods, but the reduction in activation was also correlated with self-reported nausea (Farr et al. 2016). In another study of patients with obesity and type 2 diabetes, liraglutide-induced reductions in neural response to food images were observed after short-term, but not long-term treatment (Ten Kulve et al. 2016). These correlational findings are consistent with a role for adverse effects in reduced food-cue reactivity, but the relationship among adverse effects, food-cue reactivity, and food noise requires further investigation.

Future preclinical studies using reward-magnitude-modulated PavCA could assess behavioral changes in food-cue reactivity alongside direct measures of malaise and aversion, including kaolin clay consumption and orofacial taste-reactivity responses, following acute and chronic semaglutide treatment. Measures of locomotor activity and appetite would also help distinguish malaise from general behavioral suppression or reduced food motivation. This approach would help determine whether malaise contributes to lower cue-directed responding after acute treatment and whether chronically-treated subjects continue to experience adverse effects.

One important consideration of the present study is that experiments were conducted in outbred Sprague-Dawley rats under standard feeding conditions, rather than animals with genetic or diet-induced obesity. Because GLP-1RAs are frequently prescribed for weight management in individuals with obesity, their behavioral effects may vary with body weight, metabolic state, or diet; as body weight and diet are known to influence motivated behavior. For example, rats maintained on a high-fat, high-sugar diet worked harder than control-fed rats to obtain sucrose pellets under a progressive ratio schedule of reinforcement (la Fleur et al. 2007). A high-fat diet can also alter the time course of GLP-1RA effects on feeding behavior in rats, as demonstrated with exendin-4 and liraglutide (Mul et al. 2013). However, chronic semaglutide enhanced licking for sucrose in a brief-access taste-reactivity test in mice with diet-induced obesity (Acosta et al. 2026), suggesting that enhanced responding to palatable food following extended treatment may occur across metabolic states. Direct comparisons of GLP-1RA effects on motivated behavior in standard-fed rodents and rodents with diet-induced or genetic obesity are needed to test this possibility. Nevertheless, examining GLP-1RA effects in animals without obesity remains important, given the increasing interest in these drugs as potential treatments for non-obesity related conditions, such as substance use disorders and other psychiatric conditions (Meshkat et al. 2025).

The effects of chronic semaglutide on palatable food motivation may depend on reward availability and the degree of satiation during testing. In the present study, forty 45-mg reward pellets were delivered over approximately 33 min in each behavioral session, which may not have been sufficient to induce satiation. In longer PavCA sessions involving more trials or pellets, chronically treated rats may reach satiation sooner than vehicle-treated rats and consequently exhibit less sign-tracking as the motivational value of the outcome declines.

Consistent with this possibility, chronic semaglutide reduces the time to meal cessation in rats (Cawthon et al. 2023) and decreases consumption of freely available reward pellets (Chang et al. 2026). Manipulating reward availability, session duration, and satiety state would help determine how the effects of semaglutide on cue motivation interact with its satiating effects. In a subset of rats, extending a session from 20 to 40 trials did not produce detectable treatment-related differences in behavioral responding or pellet consumption (data not shown). However, this exploratory observation may have been limited by sample size, and longer sessions or more direct manipulations of satiety may be necessary to detect treatment-dependent satiation effects.

Another limitation of the present study is that chronic treatment was administered before and during PavCA acquisition, whereas acute treatment occurred after cue-reward learning. Thus, chronic semaglutide could affect both the acquisition and expression of conditioned responding, whereas acute semaglutide was tested only for its effect on the expression of previously acquired behavior. The regimens also differed in cumulative drug exposure, dose-escalation history, and potential tolerance to adverse effects. Consequently, the experiment demonstrates differences between treatment regimens but does not isolate treatment duration as the source of those differences. Chang et al. (2026) found that administering chronic semaglutide both before and after initial PavCA training enhanced responding for access to the reward-paired cue, suggesting that chronic treatment can influence both cue-reward learning and cue motivation after learning. Nevertheless, future studies should compare acute and chronic treatments during equivalent phases of acquisition or expression of Pavlovian conditioned approach behavior.

Finally, although rats in the present study underwent dose escalation and repeated treatment for several weeks, humans may take GLP-1RAs for months or years. Future experiments could therefore extend ‘chronic’ semaglutide treatment over longer periods to determine whether changes in cue-motivated behavior persist, diminish, or reverse over timescales more comparable to clinical use. Cross-species differences in GLP-1RA pharmacokinetics must also be considered. Semaglutide has a much shorter half-life in rats than in humans, where its prolonged half-life permits once-weekly administration (Lau et al. 2015). Thus, future translational studies should consider species-specific drug exposure rather than relying solely on treatment duration or dosing frequency.

In summary, the present study demonstrates that chronic and acute semaglutide treatment regimens are associated with opposing patterns of cue-motivated behavior in a reward-magnitude-modulated PavCA paradigm. Chronic treatment increased cue-directed responding, whereas a single semaglutide dose was associated with lower sign-tracking, decreased pellet consumption, and less food-cup responding during the non-cue periods. More research is needed to determine whether these behavioral effects reflect adverse effects associated with initial drug exposure, adaptations to chronic treatment, differences in cue-reward learning phase, or other biological mechanisms. Future preclinical studies examining the effects of GLP-1RAs should carefully consider the method and schedule of drug administration and use behavioral measures that distinguish changes in incentive motivation from malaise, aversion, satiation, and nonspecific behavioral suppression. Whether these regimen-dependent effects generalize to maladaptive reward seeking or clinical outcomes remains unknown.

Determining the neural and behavioral mechanisms underlying GLP-1RA-associated changes in reward-driven behavior across stages of treatment will be important when evaluating these drugs for novel therapeutic applications beyond obesity.

## Author Note

### CRediT Statement

SSD-conceptualization, methodology, investigation, writing-original draft preparation; RL-investigation, writing-review and editing; SBF-supervision, funding acquisition, writing-review and editing.

## Acknowledgments

The authors would like to thank Lea Hsu and Juliet Franklin for their assistance with conducting the behavioral experiments and members of the Flagel Laboratory for their support and insight regarding this research.

