## Supplemental Materials for "Pavlovian conditioned approach with different reward magnitudes reveals opposing effects of acute and chronic semaglutide treatment"

**Table S1.** *ANOVA statistical results for sessions 6-7 reward-magnitude-modulated PavCA behavioral data for vehicle and acute semaglutide treatment groups shown in **Fig. S1**.*

Significant effects ($p < 0.05$) are bolded and italicized.

| | Lever Contacts | Probability of Lever Contact | Latency to Lever Contact | Food Cup Entries | Probability of Food Cup Entry | Latency to Enter Food Cup |
|---|---|---|---|---|---|---|
| Sex | F(1,38)= 1.120, p=0.297 | F(1,38)= 1.260, p=0.269 | F(1,38)= 3.120, p=0.085 | F(1,38)= 2.595, p=0.115 | F(1,38)= 1.775, p=0.191 | F(1,38)= 2.851, p=0.100 |
| Treatment | ***F(1,38)= 6.272, p=0.017*** | ***F(1,38)= 14.249, p<0.001*** | ***F(1,38)= 12.711, p=0.001*** | F(1,38)= 2.019, p=0.164 | F(1,38)= 2.151, p=0.151 | F(1,38)= 2.184, p=0.148 |
| Reward Magnitude | ***F(1,38)= 11.659, p=0.002*** | ***F(1,38)= 7.140, p=0.011*** | ***F(1,38)= 15.598, p<0.001*** | F(1,38)= 0.529, p=0.471 | F(1,38)= 0.062, p=0.803 | F(1,38)= 3.157, p=0.084 |
| Session | ***F(1,38)= 32.229, p<0.001*** | ***F(1,38)= 90.321, p<0.001*** | ***F(1,38)= 57.638, p<0.001*** | ***F(1,38)= 17.374, p<0.001*** | ***F(1,38)= 20.426, p<0.001*** | ***F(1,38)= 16.970, p<0.001*** |
| Sex* Treatment | F(1,38)= 0.037, p=0.848 | F(1,38)= 0.137, p=0.713 | F(1,38)= 0.076, p=0.784 | F(1,38)= 0.158, p=0.693 | F(1,38)= 0.006, p=0.939 | F(1,38)= 0.084, p=0.773 |
| Sex* Magnitude | F(1,38)= 0.298, p=0.589 | F(1,38)= 0.487, p=0.490 | F(1,38)= 0.004, p=0.949 | F(1,38)= 0.000, p=0.992 | F(1,38)= 0.038, p=0.847 | F(1,38)= 0.136, p=0.714 |
| Sex*Session | F(1,38)= 0.495, p=0.486 | F(1,38)= 1.585, p=0.216 | F(1,38)= 0.864, p=0.358 | F(1,38)= 0.132, p=0.718 | F(1,38)= 0.107, p=0.746 | F(1,38)= 0.109, p=0.743 |
| Treatment* Magnitude | F(1,38)= 2.361, p=0.133 | F(1,38)= 1.004, p=0.323 | F(1,38)= 2.687, p=0.109 | F(1,38)= 1.281, p=0.265 | F(1,38)= 1.014, p=0.320 | F(1,38)= 0.192, p=0.664 |
| Treatment* Session | ***F(1,38)= 53.349, p<0.001*** | ***F(1,38)= 115.632, p<0.001*** | ***F(1,38)= 97.788, p<0.001*** | ***F(1,38)= 7.195, p=0.011*** | F(1,38)= 2.367, p=0.132 | F(1,38)= 2.278, p=0.139 |
| Magnitude *Session | ***F(1,38)= 9.127, p=0.004*** | F(1,38)= 0.308, p=0.582 | F(1,38)= 3.776, p=0.059 | F(1,38)= 0.937, p=0.339 | F(1,38)= 2.874, p=0.098 | F(1,38)= 0.108, p=0.745 |
| Sex* Treatment* Magnitude | F(1,38)= 0.014, p=0.908 | F(1,38)= 1.191, p=0.282 | F(1,38)= 0.042, p=0.839 | F(1,38)= 0.000, p=0.992 | F(1,38)= 0.095, p=0.759 | F(1,38)= 0.101, p=0.752 |
| Sex* Treatment* Session | F(1,38)= 2.240, p=0.143 | F(1,38)= 0.402, p=0.530 | F(1,38)= 2.853, p=0.099 | F(1,38)= 1.470, p=0.233 | F(1,38)= 2.567, p=0.117 | F(1,38)= 1.438, p=0.238 |
| Sex* Magnitude* Session | F(1,38)= 0.030, p=0.864 | F(1,38)= 0.922, p=0.343 | F(1,38)= 0.004, p=0.951 | F(1,38)= 0.105, p=0.748 | F(1,38)= 0.604, p=0.442 | F(1,38)= 0.433, p=0.514 |
| Treatment* Magnitude* Session | F(1,38)= 0.018, p=0.893 | F(1,38)= 2.192, p=0.147 | F(1,38)= 0.120, p=0.731 | F(1,38)= 1.710, p=0.199 | F(1,38)= 0.435, p=0.514 | F(1,38)= 0.169, p=0.683 |
| Sex* Treatment* Magnitude* Session | F(1,38)= 0.503, p=0.483 | F(1,38)= 0.039, p=0.845 | F(1,38)= 0.022, p=0.883 | F(1,38)= 0.193, p=0.663 | F(1,38)= 0.604, p=0.442 | F(1,38)= 0.736, p=0.396 |

**Table S2.** *ANOVA (pretraining/session 1; PavCA session 7) and LMM (sessions 1-6) statistical results for pellets remaining and non-CS food cup entry data shown in **Fig. S2**.* Significant effects (p < 0.05) are bolded and italicized.

| | Pretraining/ Session 1 Pellets Remaining (Prop.) | Pretraining/ Session 1 Non-CS Food Cup Entries / minute | Sessions 1-6 Pellets Remaining (Prop.) | Sessions 1-6 Non-CS Food Cup Entries / minute | Session 7 Pellets Remaining (Prop.) | Session 7 Non-CS Food Cup Entries / minute |
|---|---|---|---|---|---|---|
| Sex | F(1,60)= 1.812, p=0.183 | F(1,60)= 0.488, p=0.488 | F(1,61.067)= 0.822, p=0.368 | F(1,70.218)= 0.015, p=0.904 | F(1,26)= 0.751, p=0.394 | F(1,26)= 0.013, p=0.910 |
| Treatment | F(1,60)= 2.935, p=0.092 | ***F(1,60)= 28.235, p<0.001*** | F(1,61.067)= 0.957, p=0.332 | F(1,70.218)= 3.272, p=0.075 | ***F(2,26)= 13.812, p<0.001*** | ***F(2,26)= 3.631, p=0.041*** |
| Session | F(1,60)= 0.682, p=0.412 | ***F(1,60)= 5.649, p=0.021*** | F(5,134.725)= 0.475, p=0.795 | ***F(5,99.932)= 20.985, p<0.001*** | NA | NA |
| Sex* Treatment | F(1,60)= 1.812, p=0.183 | F(1,60)= 0.649, p=0.424 | F(1,61.067)= 0.822, p=0.368 | F(1,70.218)= 0.059, p=0.808 | F(2,26)= 0.763, p=0.476 | F(2,26)= 0.105, p=0.901 |
| Sex*Session | F(1,60)= 0.153, p=0.697 | F(1,60)= 0.135, p=0.715 | F(5,134.725)= 0.620, p=0.684 | F(5,99.932)= 1.743, p=0.132 | NA | NA |
| Treatment* Session | F(1,60)= 0.682, p=0.412 | F(1,60)= 3.428, p=0.069 | F(5,134.725)= 0.475, p=0.795 | ***F(5,99.932)= 3.905, p=0.003*** | NA | NA |
| Session*Sex* Treatment | F(1,60)= 0.153, p=0.697 | F(1,60)= 1.888, p=0.174 | F(5,134.725)= 0.620, p=0.684 | F(5,99.932)= 0.326, p=0.896 | NA | NA |

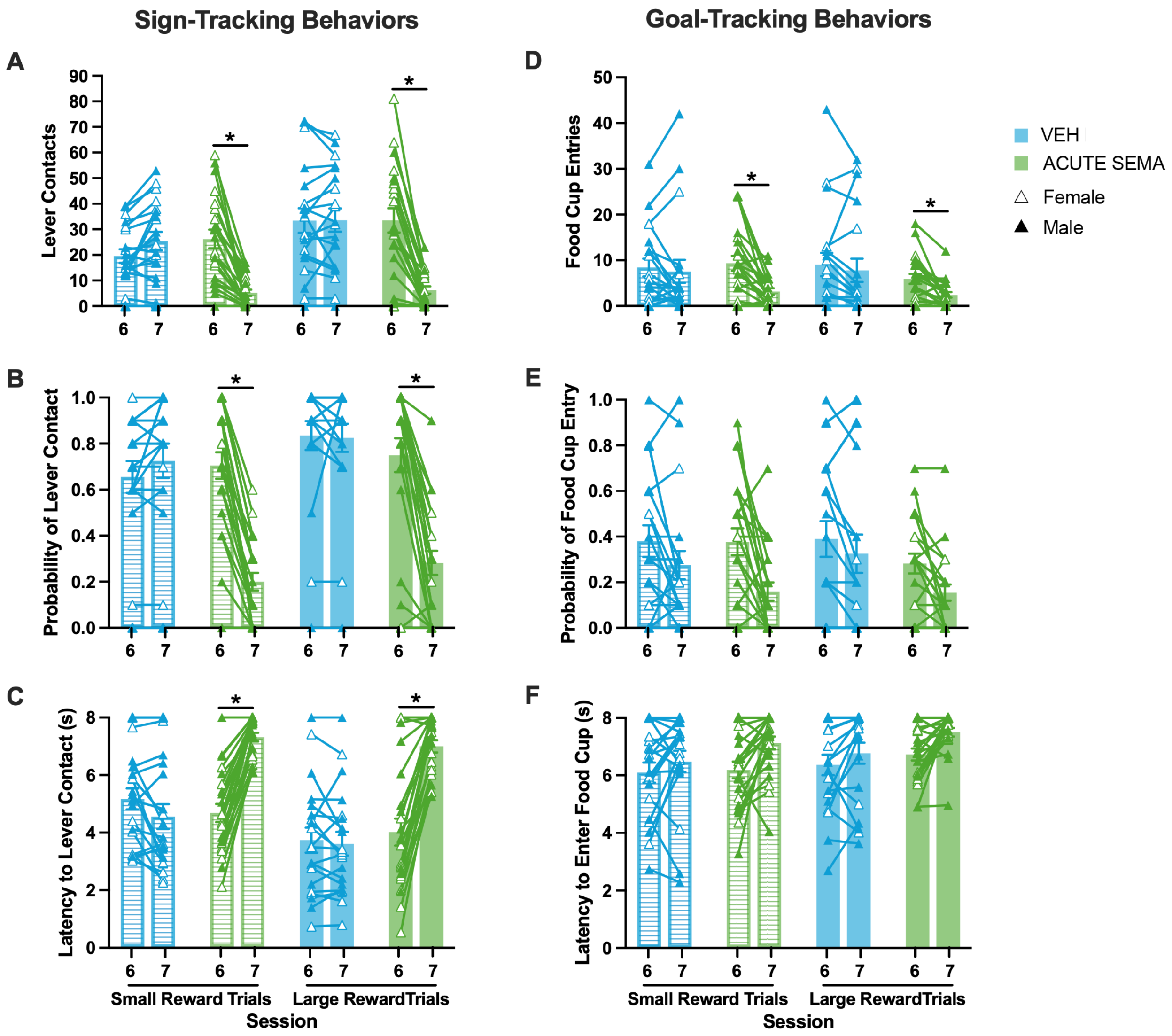


**Fig. S1** *Acute semaglutide treatment reduces Pavlovian conditioned approach behavior.* Bar graphs behavioral measures for small and large reward trials (mean +/- SEM; triangular points represent individual subjects) over sessions 6-7 of reward-magnitude-modulated PavCA (RMM-PavCA) for the vehicle and acute semaglutide treatment groups. Lever-cue directed (i.e. sign-tracking) behavioral measures included **(A)** contacts, **(B)** probability of contact, and **(C)** latency to contact the lever-cue. Food cup directed (i.e. goal-tracking) behaviors included **(D)** entries, **(E)** probability of entry, and **(F)** latency to enter the food cup. *Indicates $p < 0.05$ for post-hoc session comparison within treatment group.

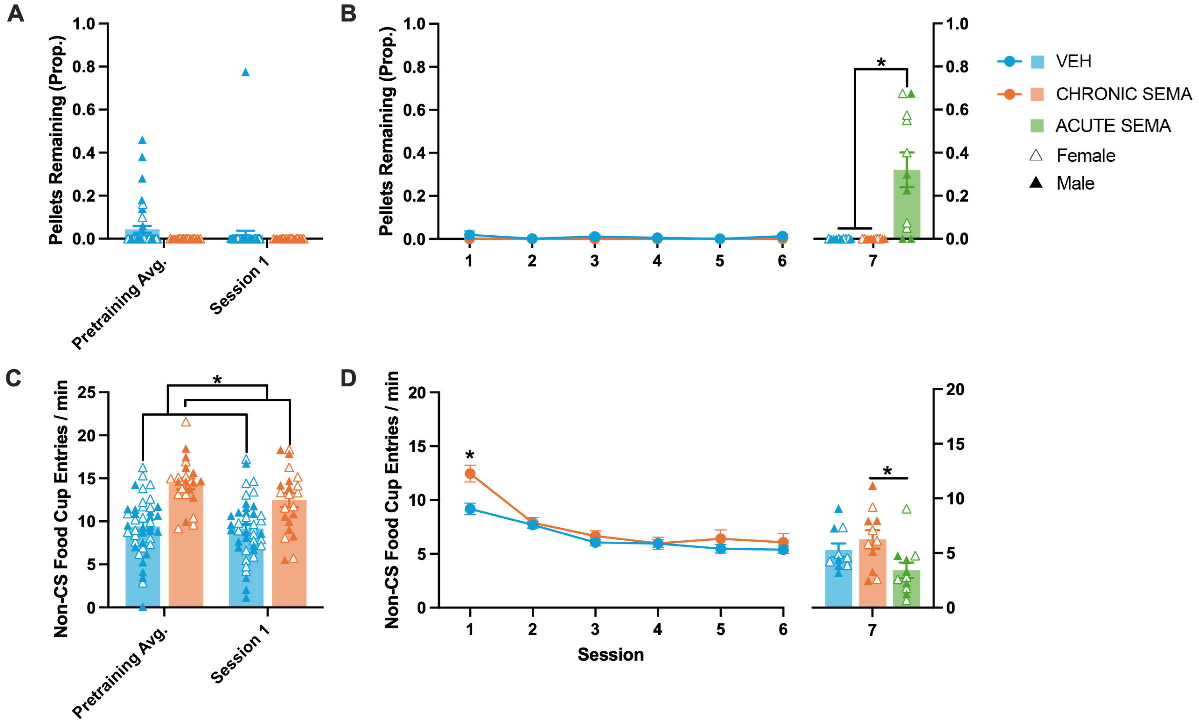


**Fig. S2** *Chronic and acute semaglutide treatment have distinct effects on reward-pellet consumption and non-CS food-cup entries during behavioral testing.* Graphs for **(A-B)** the number of pellets remaining of the total delivered and **(C-D)** the number of food cup entries per minute during the non-CS periods. **(A,C)** Bar graphs show data (mean +/- SEM; triangular points represent individual subjects) across pretraining (averaged over 2 sessions) and reward-magnitude-modulated PavCA (RMM-PavCA) session 1, for chronic semaglutide and vehicle treatment groups (*indicates $p < 0.05$ for main effect of treatment). **(B,D)** Line graphs show data (mean +/- SEM) across sessions 1-6 of RMM-PavCA, for chronic semaglutide and vehicle treatment (*indicates $p < 0.05$ for post-hoc treatment comparison within session). Bar graphs show the same measures (mean +/- SEM; triangular points represent individual subjects) on session 7 of RMM-PavCA, including the acute semaglutide treatment group (*indicates $p < 0.05$ for post-hoc treatment comparison).
